# Mitotic catastrophe and other cellular instability events in sodium valproate-treated HeLa cells

**DOI:** 10.64898/2026.08.22.746408

**Authors:** Beatriz Pierre Sforça, Camila Borges Oliveira, Mariana Minski Furtado, Milena Gomes Santos, Marina Amorim Rocha, Maria Luiza S. Mello

**Affiliations:** Department of Structural and Functional Biology, Institute of Biology, University of Campinas (UNICAMP), 13083-862 Campinas, SP, Brazil

**Author notes:** These authors share first authorship (Beatriz Pierre Sforça, Camila Borges Oliveira). Corresponding author. *E-mail address* (M.L.S. Mello).

**Keywords:** Cell death, Valproic acid, HeLa cells, Caspase-2, p53, DNMTs

## Abstract

Valproic acid/sodium valproate (VPA) is a widely prescribed anticonvulsant and has also been used against certain tumor cells. It is a potent modulator of gene expression. Its ability to induce apoptosis has been well documented in HeLa cells. However, another form of cell death - mitotic catastrophe - has not yet been explored in VPA-treated HeLa cells. Here, we investigated the effects of VPA treatment on mitotic catastrophe characteristics, including morphological features and their frequencies, fluorescence intensity signals of caspase-2 and p53, and the expression and abundance of DNMT1 and DNMT3B. An increased frequency of mitotic catastrophe was observed not only morphologically, but also through enhanced induction of caspase-2, involvement of p53, at least under more drastic VPA treatment, but without a decrease in DNMT1 or DNMT3B levels. Additionally, enhancement of mitotic catastrophe coincided with a reduction in mitotic chromosome abnormalities. Increased *DNMT3B* expression following VPA action, may be favored by previously reported chromatin decondensation induced by this drug. Enhanced CpG methylation of specific DNA sites could thus be promoted. In conclusion, VPA was shown to trigger metabolic pathways linked to different forms of cell death in HeLa cells, supporting its oncosuppressive potential.

## 1. Introduction

Valproic acid associated with its sodium salt (VPA), a drug widely prescribed as an anticonvulsant, is a gene expression modulator that inhibits histone deacetylases, affects the DNA and histone methylation status, induces structural and functional chromatin remodeling, and may interact directly with DNA and histones (Göttlicher et al., 2001; Phiel et al., 2001; Eyal et al., 2005; Sargolzaei et al., 2017; Veronezi et al., 2017; Romoli et al., 2019; Rocha et al., 2019, 2023; Vidal and Mello, 2020; Mello, 2021).

In HeLa cells, a cell lineage originating from human cervical cancer, VPA induces DNA demethylation through a predominantly active process involving enzymes of the “ten-eleven translocation” (TET) family and thymine DNA glycosylase (TDG) (Rocha et al., 2019). In addition, VPA induces a passive DNA demethylation process that affects the levels of DNA-methyltransferase 1 (DNMT1) (Rocha et al., 2019), an enzyme required for the maintenance of overall methylation levels in the genome (Zhang and Xu, 2017; Lyko, 2018). DNMT1 is overexpressed in solid tumors and affects the activity of several tumor suppressor genes in cervical tumorigenesis (Saito et al., 2003; Peng et al., 2006; Zhang et al., 2011; Zhang and Xu, 2017; Lyko, 2018).

Cell cycle arrest and apoptosis via intrinsic and extrinsic pathways are induced in HeLa cells cultured in the presence of VPA (Dejligbjerg et al., 2008; Han et al., 2013; Hashemi et al., 2022; Kar et al., 2026), although no change in cell death was reported under VPA doses close to those reached in the plasma after therapeutic antiepileptic purposes (0.3 mM-0.7 mM) (Felisbino et al., 2011). Activation of caspases-3, -8, and -9 and loss of mitochondrial membrane potential in HeLa cells treated with 10 mM VPA for 24 h have been demonstrated statistically (Han et al., 2013). An increase in the expression of pro-apoptotic genes, including *p21* and *p53*, as well as reduction in the expression of *Bcl-2*, an anti-apoptotic gene, have also been reported for HeLa cells treated with VPA for 48 h at concentrations higher than 25 mM; under this experimental condition, the percentages of late apoptosis were 17.10% as compared to 1.31% in the control group (Hashemi et al., 2022). An increase in *p53* expression has also been reported for HeLa cells cultured in the presence of 25 and 50 mM VPA for 24 h (Kar et al., 2026).

Despite the reports on induction of apoptosis in HeLa cells after VPA treatment, mitotic catastrophe, a form of cell death preceded by multi-nucleation linked to delayed mitosis (Galluzzi et al., 2012), and firstly described in this cell line (Swanson et al., 1995; Miranda et al., 1996) has not been explored, except by a report of cells containing multinucleated bodies in non-synchronized cell preparations (Felisbino et al., 2011). Mitotic catastrophe is considered an onco-suppressive pathway preceding apoptosis, necrosis, or senescence through which genomic instability may be avoided and mitosis-incompetent cells are eliminated (Vitale et al., 2011; Galluzzi et al., 2012; Sazonova et al., 2021). Although caspase-2 and p53 proteins have been associated to the incidence of mitotic catastrophes (Castedo et al., 2004; Vitale et al. 2017; Sazonova et al. 2021), and a relationship between DNMT1 depletion and increased incidence of mitotic catastrophe has been reported in human solid tumor cells (Chen et al., 2007), this association has not been studied in HeLa cells cultured in the presence of VPA.

In the present study, we investigated the morphological characteristics and frequency of mitotic catastrophe, and other cellular instability events, such as mitotic defects, in synchronized HeLa cells cultured with varying concentrations of VPA over different exposure times. We also analyzed the incidence of caspase-2 and p53 fluorescence signals, along with gene expression and the abundance of DNMT1. In addition, gene expression and abundance of DNMT3B, which are frequently overexpressed in HeLa cells and that are linked to mitotic chromosome condensation, playing a role in epigenetic regulation and genome stability (Geiman et al., 2004), were also investigated. This study aimed to contribute to expanding our knowledge of the multitarget effects of VPA.

## 2. Materials and Methods

### 2.1. Cell line and culture conditions

HeLa cells (ATCC:CCL-2) acquired at passage ten from the Emerging Virus Studies Laboratory at UNICAMP (Campinas, Brazil) and validated at the Technical Directorate for Teaching and Research Support of the University of São Paulo (São Paulo, Brazil) were used at passages 14-22 and cultured at 1.5 x 10^5^ cells mL^-1^, in 6-well plates, in high-glucose Dulbecco’s modified Eagle’s medium (Sigma-Aldrich®; Merck KGaA) supplemented with 10% fetal calf serum (FCS; Nutricell Cellular Nutrients, Campinas, SP, Brazil), penicillin-streptomycin (Sigma-Aldrich®, St. Louis, MO, USA – 100 IU mL^-1^ and 100 µg mL^-1^, respectively), and 1% sodium pyruvate (Sigma-Aldrich®; Merck KGaA) at 37 °C in 5% CO_2_ atmosphere.

Cell synchronization was performed as previously reported (Rocha et al. 2019). Briefly, to obtain cells arrested in the G1 phase at the optimal cell concentration, the cells were initially plated and cultured in 1% FCS medium containing 20 µM lovastatin (Sigma-Aldrich) for 24 h. The preparations were then washed with phosphate-buffered saline (PBS) and cells were induced to continue cycling by replacing the medium with 10% FCS containing 6 mM mevalonic acid (Santa Cruz Biotechnology, Dallas, TX, USA) for 18 h to obtain cells in the S phase. No change in viability has been reported for HeLa cells under the synchronization procedure that uses the 3-(4,5-dimethylthiazol-2-yl)-2,5-diphenyltetrazolium bromide (MTT) assay (Rocha et al., 2019, 2021).

### 2.2. Cell treatment designed for analysis of chromosome abnormalities and cell death morphological identification

HeLa cells were cultured in medium containing 1% FCS and 1 or 20 mM VPA (Sigma-Aldrich) for 4 h and 5 mM VPA for 48 h, preceded by culture for 24 h in the absence of the drug. VPA concentrations and treatment time were selected based on cell viability as detected with the MTT assay. Cells cultured for 4 h in 1 or 20 mM VPA exhibited viability levels of 100% and 98%, respectively (Rocha et al. 2019, 2021). Cells showed less than 50% of viability when cultured in 5 mM VPA for 48 h. Controls consisted in cells cultured in the absence of the drug.

Cells were fixed in an absolute ethanol-glacial acetic acid mixture (3:1, v/v) for 1 min, rinsed with absolute ethanol, and air-dried. They were then subjected to the Feulgen reaction (Mello and Vidal, 2017), in which treatment with 4 M HCl for 1 h at 25 °C was used as the hydrolysis step. After treatment with Schiff reagent for 40 min, the preparations were rinsed thrice in a bath of sulfurous water (5 min each time) and one bath of distilled water, air-dried, and rapidly counterstained with fast green at pH 2.7 for improving identification of cell individuality. The preparations were then rinsed with distilled water, air-dried, cleared in xylene for 10 min, mounted on Canada balsam, and examined using a Nikon light microscope (Tokyo, Japan) for detection of the frequencies of typical images of mitotic catastrophes, apoptosis, cells exhibiting mitotic defects, and single giant nuclei. These frequencies were established for randomly selected 2000 cells for three preparations of each cell treatment and control. Images were captured using an Axiophot microscope equipped with an AxioCamHRc camera (Carl Zeiss, Oberkochen, Germany). Cells exhibiting multinucleation and membrane-involved lightly stained nuclear fragments were considered to have undergone mitotic catastrophe. Images in which deeply stained vesicles are discriminated, indicated apoptosis. Mitotic abnormalities included tri- and tetrapolar metaphases, metaphases and anaphases containing lagging chromosome fragments, and anaphases, telophases, and cytokinesis exhibiting chromosome bridging. The frequencies of these features per 2000 cells per preparation were determined.

### 2.3. Caspase-2 and p53 immunocytochemistry

HeLa cells treated with 1 mM or 20 mM VPA for 4 h, as well as those treated with 20 mM VPA for 48 h, were fixed in methanol at -20 °C for 10 min and subsequently rinsed once in PBS. All cells were cultured for 24 h without the drug prior to fixation. Given the association of caspase-2 and p53 proteins with the occurrence of mitotic catastrophes (Vitale et al. 2017; Sazonova et al. 2 021), we employed an immunofluorescence protocol based on previously conducted tests to detect these proteins. The cells were permeabilized using 0.2% Triton X-100 (cat. no. T-8787; CAS no. 9002-93-1; Merck, Darmstadt, Germany) for 10 min at 25 °C, and then blocked with 5% BSA for 1 h at the same temperature. The preparations were then incubated overnight at 4 °C with rabbit anti-caspase-2 recombinant primary monoclonal antibody (Invitrogen, Thermo-Fisher Sci., Waltham, MA, USA) (Lot no. YA37955291; dilution 1:200 in 1% BSA) and rabbit anti-p53 primary polyclonal antibody (Rheabiotech, Campinas, SP, Brazil) (Lot no. 23080; dilution 1:100 in 1% BSA), rinsed thrice in PBS (pH 7.4; 5 min each), followed by incubation at 25 °C for 2 h with goat anti-rabbit secondary antibody conjugated with Alexa-Fluor 488 fluorochrome (Life Technologies, Carlsbad, CA, USA) (no. A-11,008, Lot no. 2284,594; dilution 1:500 in 1% BSA). The preparations were then rinsed in PBS and mounted in VECTASHIELD® (cat. no. H-1000, Lot no. V1001; Vector Laboratories, Inc., Burlingame, CA, USA). For control cells cultured for 4 h, the assays were repeated three times, two of each performed in duplicate (technical replicates). For cells cultured for 4 h in the presence of 1 mM VPA, the assays were repeated five times and with technical replicates. For cells cultured for 4 h in the presence of 20 mM VPA, the assays were repeated three times and with technical replicates. For cells cultured with or without 5 mM VPA for 48 h, the assays were repeated three times; however, biological replicates were not available in this case.

Fluorescence signals for caspase-2 and p53 were captured using a Leica TCS SS ll (Wetzlar, Germany) confocal microscope at the LaCTAD facilities (UNICAMP, Campinas, Brazil) and analyzed using ImageJ software (NIH, USA).

A dataset containing all fluorescence signal values was deposited in the data and metadata repository of the University of Campinas (UNICAMP, Campinas, Brazil) (Sforça and Mello, 2026).

### 2.4. RT-qPCR

Cells designed for the analysis of the expression of the *DNMT1* and *DNMT3B* genes were treated with 1 mM VPA for 4 h or with 2 mM VPA for 48 h, and respective controls, following cell culture for 24 h in the absence of the drug. They were subjected to total RNA isolation using RNeasy Mini kit (Qiagen®, Hilden, Germany) according to the manufacturer’s instructions. RNA integrity number of the samples was evaluated using a Nanodrop™ spectrophotometer (Romford, England). Two µg RNA per sample were employed for cDNA synthesis using the High-Capacity cDNA Reverse Transcription kit (Applied Biosystems™, Chicago, IL, USA). The TaqMan probes specific for *DNMT1*, *DNMT3B*, and the housekeeping gene *GAPDH* were as follows: Hs00945875_m1, Hs01003405_m1, and Hs02758991_g1, respectively. The PCR primers were not provided by the drug manufacturer.

Cycle threshold values were calculated from experiments performed in triplicate for *DNMT1* and in five times for *DNMT3B*, and normalized with respect to the housekeeping gene *GAPDH*. PCR was conducted on an Applied Biosystems 7500 real-time PCR system (Applied Biosystems®) and relative quantification of *DNMT1* and *DNMT3B* was achieved using the comparative 2^-ΔΔCq^ method (Livak and Schmittgen 2001).

### 2.5. Western blotting

Cells designed for the analysis of the DNMT1 and DNMT3B protein abundance were treated with 1 mM VPA for 4 h and with 2 mM VPA for 48 h, as well as respective controls, following cell culture for 24 h in the absence of the drug. Total proteins were extracted using RIPA buffer (50 mM Tris-HCl at pH 8.0, 150 mM NaCl, 1% Triton X-100, 0.5% sodium deoxycholate, 0.1% SDS, 1 mM EDTA, 0.5 mM EGTA, and 1 mM PMSF) for 30 min on ice, followed by centrifugation at 14,000 RCF for 15 min. The Bradford assay (Sigma-Aldrich®) was used to detect protein concentrations, using BSA as a standard. Absorbance values were obtained after the samples were incubated for 1 h at room temperature at λ = 595 nm, using a Multiskan™ FC Microplate Photometer (Thermo Fisher Scientific, Inc.). Protein samples (30 µg) were incubated in heated sample buffer (0.06 M Tris-HCl at pH 6.8, 2% SDS, 10% glycerol, 5% β-mercaptoethanol, 0.025% Bromophenol Blue) for 5 min and separated by SDS-PAGE on 17% polyacrylamide gels. The proteins were transferred to nitrocellulose membranes (Thermo Fisher Scientific, Inc.) which were blocked in 4% BSA at 25 °C for 2 h, and separately incubated with rabbit anti-DNMT1 (Cell Signaling Technol., Danvers, USA; dilution 1:1,000) and rabbit anti-DNMT3B (Cell Signaling Technol.; dilution 1:1,000) primary antibodies overnight in 1 x Tris-buffered saline-0.1% Tween 20 (TBST; cat. no.91414; Sigma-Aldrich®) blocking solution at 4 °C. After extensive washing with TBST, the membranes were incubated with horseradish peroxidase-conjugated goat anti-rabbit (Chemicon®, Billerica, MA, USA; dilution: 1:5,000) secondary antibody for 2 h at 25 °C in 1% BSA blocking solution followed by extensive washes. Protein blots were imaged using an ECL Western Blotting Detection System (Amersham®, Pittsburgh, PA, USA) and were detected by chemiluminescence using an Alliance 6.7 (UVITEC, Cambridge, UK) image system at the Obesity and Comorbidities Research Center (Institute of Biology, UNICAMP). The assays were conducted using three different cell passages for DNMT1 and four different cell passages for DNMT3B.

### 2.6. Statistics

Statistics was performed using GraphPad Prism version 8.0.1 (Boston, MA, USA). Shapiro-Wilk test was used to detect the data distribution character. Student’s t-test and Mann-Whitney test were used to assess the statistical significance of the datasets. In all cases, a p-value of <0.05 was considered the critical level for rejecting the null hypothesis.

## 3. Results

### 3.1. VPA induced increased frequency of morphological images of mitotic catastrophe followed by apoptosis, and decreased mitotic abnormalities

Images of mitotic catastrophe, typical late apoptosis, apoptosis generated from mitotic catastrophe, abnormal mitosis, and giant mononucleate nuclei, were detected in both VPA-treated and untreated HeLa cells (Fig. 1 A). The relative frequency of these images under the various VPA treatments relative to respective controls as well as the rate of their minimum-maximal absolute values and identification of the statistical test used for comparisons are shown in Fig. 1 B and Table 1, respectively.

**Fig. 1.**
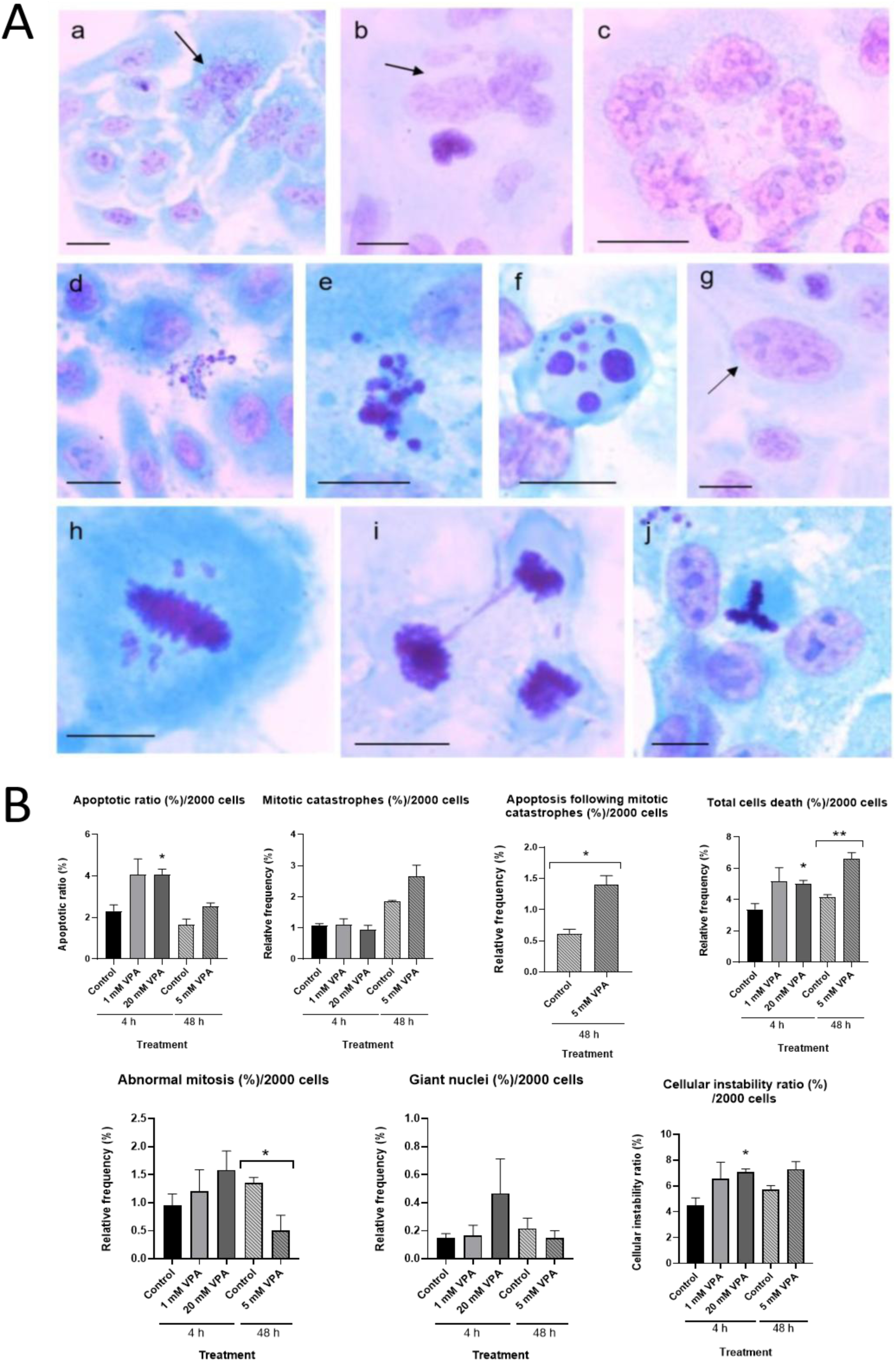
Cell death images and aspects of nuclear abnormalities in Feulgen-acid fast green-stained preparations (A) and frequency of these elements in VPA-treated and untreated HeLa cells (B). Mitotic catastrophes (a-c, arrows), typical apoptosis (d, e), apoptosis suspected to be generated from a mitotic catastrophe (f), giant nucleus (g, arrow), lagging chromosomes (h), chromosome bridging (i), and tripolar metaphase (j) (A). Scale bars, 25 µm. The vertical lines above the graphical bars in B represent the standard error of the mean. The cellular instability ratio represents the sum of the frequencies of all these elements (B). *, significant differences at the p< 0.05 level; **, significant differences at the p< 0.01 level.

**Table 1.** Cell death and cell abnormalities in VPA-treated HeLa cells.

| Treatments | Mitotic catastrophe |  |  | Apoptosis |  |  | Apoptosis following mitotic catastrophe |  |  | Total cell death |  |  |
| --- | --- | --- | --- | --- | --- | --- | --- | --- | --- | --- | --- | --- |
|  | Minimum-maximal absolute values | Frequency (%) |  | Minimum-maximal absolute values | Frequency (%) |  | Minimum-maximal absolute values | Frequency (%) |  | Minimum-Maximal absolute values | Frequency (%) |  |
|  |  | X | S |  | X | S |  | X | S |  | X | S |
| Control – 4 h | 19-24 | 1.07 | 0.12 | 40-58 | 2.32<sup>a1</sup> | 0.51 | - | - | - | 59-82 | 3.38<sup>a3</sup> | 0.63 |
| 1 mM VPA – 4 h | 16-28 | 1.12 | 0.30 | 56-108 | 4.07 | 1.30 | - | - | - | 72-131 | 3.55 | 0.58 |
| 20 mM VPA – 4 h | 15-24 | 0.95 | 0.23 | 75-91 | 4.08<sup>b1</sup> | 0.42 | - | - | - | 93-106 | 4.37<sup>b3</sup> | 0.93 |
| Control – 48 h | 36-38 | 1.85 | 0.05 | 22-40 | 1.65 | 0.48 | 11-15 | 0.62<sup>a2</sup> | 0.12 | 75-88 | 4.12<sup>a4</sup> | 0.33 |
| 5 mM VPA – 48 h | 41-65 | 2.67 | 0.60 | 44-55 | 2.53 | 0.29 | 23-33 | 1.40<sup>b2</sup> | 0.25 | 117-142 | 6.60<sup>b4</sup> | 0.66 |

| Treatments | Abnormal mitosis |  |  | Giant mononucleate cells |  |  | Total nuclear instability (NIR) |  |  |
| --- | --- | --- | --- | --- | --- | --- | --- | --- | --- |
|  | Minimum-maximal absolute values | Frequency (%) |  | Minimum-maximal absolute values | Frequency (%) |  | Minimum-maximal absolute values | Frequency (%) |  |
|  |  | X | S |  | X | S |  | X | S |
| Control – 4 h | 13-27 | 0.95 | 0.36 | 2-4 | 0.15 | 0.05 | 3.70-5.65 | 4.48<sup>a6</sup> | 1.03 |
| 1 mM VPA – 4 h | 13-39 | 1.20 | 0.67 | 1-6 | 0.17 | 0.13 | 4.30-8.80 | 6.55 | 2.25 |
| 20 mM VPA – 4 h | 23-45 | 1.58 | 0.59 | 3-7 | 0.27 | 0.10 | 6.50-7.55 | 6.88<sup>b6</sup> | 0.70 |
| Control – 48 h | 25-31 | 1.35<sup>a5</sup> | 0.48 | 2-7 | 0.22 | 0.13 | 5.10-6.20 | 5.70 | 0.53 |
| 5 mM VPA – 48 h | 3-21 | 0.50<sup>b5</sup> | 0.29 | 2-5 | 0/15 | 0.09 | 6.30-8.30 | 7.30 | 1.00 |

Analysis done in triplicate; n = 2000 cells. Different letters in each column indicate significant differences at the p<0.05 level (a1, b1; a3, b3; a6, b6 – ANOVA) (a2, b2; a4, b4; a5, b5 – t-test). S, standard deviation; X, arithmetic mean. The individual values composed a dataset that was deposited in a public repository at the University of Campinas (UNICAMP) (Sforça et al. 2026).

The frequency of mitotic catastrophes, when identified by the presence of less packed multinucleate bodies inside cells (Fig. 1 Aa-c), were not affected by the VPA treatments (Fig. 1 B). However, if images of deeply stained multinucleate bodies are observed inside the cellular structures are considered in separate, suggesting apoptosis following mitotic catastrophe (Fig. 1 Af), a significant increase of the frequency of this parameter was induced by a 48-h-treatment with 5 mM VPA (Fig. 1 B). Late apoptotic images properly, in which apoptotic vesicles are observed free from cellular contours (Fig. 1 Ad,e), had their frequency increased after a 4-h-treatment with 20 mM VPA (Fig. 1B). Summing these forms of cell death into one parameter revealed significant increases after treatments with 20 mM VPA for 4 h and 5 mM VPA for 48 h (Fig. 1 B). The frequency of the giant mononucleate nuclei was not affected by VPA treatments (Fig. 1 Ag, B).

Abnormal mitoses, including cells exhibiting lagging chromosomes, chromosome bridging, and tri- and tetrapolar metaphases (Fig. 1 Ah-j) had their frequencies decreased under the 48-h cell treatment with 5 mM VPA (Fig. 1 B). All these data put together resulted in a cellular instability ratio which differed only for cells cultured in the presence of 20 mM VPA for 4 h (Fig. 1 B).

### 3.2. VPA induced increasing caspase-2 marks but only a trend to increasing p53 marks

A significant increase in the intensity of the fluorescence signals for caspase-2 occurred in HeLa cells treated with 20 mM VPA for 4 or 48 h relative to untreated controls (Fig. 2 A, B). Cells treated with 1 mM VPA for 4 h did not show differences when compared to untreated control (Fig. 2 B).

**Fig. 2.**
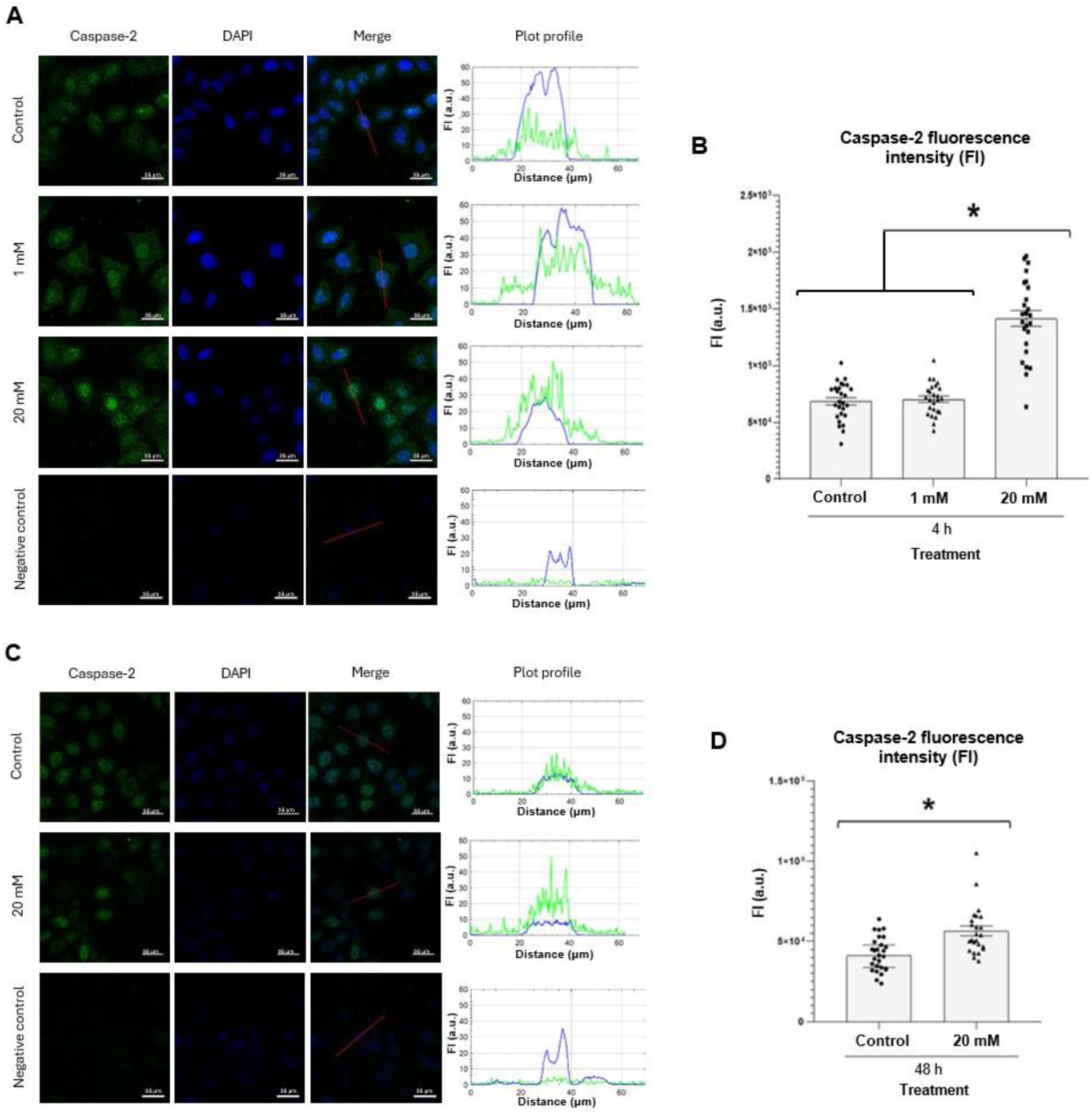
Immunofluorescence intensity of caspase-2 signals in VPA-treated HeLa cells, as assessed using confocal microscopy. Images A and C are representative of independent experiments as mentioned in the Materials and Methods section. The total number of nuclei analyzed is detailed elsewhere (Sforça and Mello, 2026). Scale bars indicate 25 μm. Graphs represent fluorescence intensity profiles along the red line drawn in the merged image of selected nuclear images to identify the immunofluorescence signals for caspase-2 (green) and DAPI-stained DNA (blue). Fluorescence intensity of caspase-2 signals increased in response to 20 mM VPA treatments for 4 h and 48 h relative to untreated controls (B, D). *, significant difference at p*<* 0.05 level (Mann-Whitney test). The vertical lines above the graphical bars represent the standard error of the mean. FI, fluorescence intensity in arbitrary units (a.u.).

No significant changes were detected in the intensity of the fluorescence signals for p53 between VPA-treated and untreated cells, although a trend to increased values is suggested for cells treated with 20 mM VPA for 48 h (Fig. 3 A, B).

**Fig. 3.**
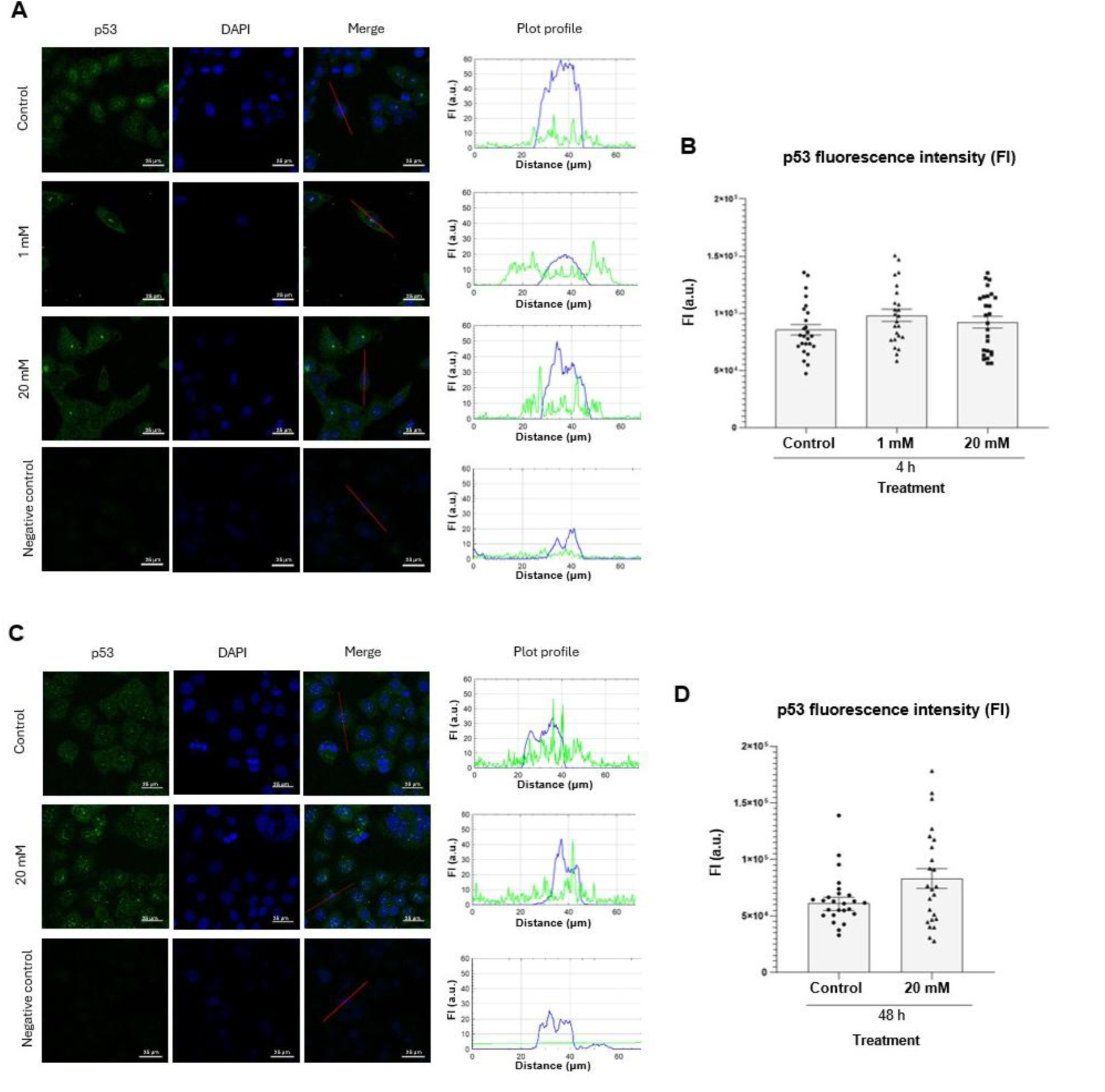
Immunofluorescence intensity of p53 signals in VPA-treated HeLa cells, as assessed using confocal microscopy. Images A and C are representative of independent experiments as mentioned in the Materials and Methods section. The total number of nuclei analyzed is detailed elsewhere (Sforça and Mello 2026). Bars = 25 μm. Graphs represent fluorescence intensity profiles along the red line drawn in the merged image of selected nuclear images to identify the immunofluorescence signals for p53 (green) and DAPI-stained DNA (blue). Significant differences in fluorescence signals between VPA-treated and untreated cells were not detected at the p < 0.05 level (B, D). The vertical lines above the graphical bars represent the standard error of the mean. FI, fluorescence intensity in arbitrary units (a.u.).

### 3.3. VPA induced a significant increase in DNMT3B expression and a trend to increase in DNMT1 expression and DNMT3B abundance

There was no significant alteration in mRNA expression levels of *DNMT1* after cell treatment with 1 mM VPA for 4 h and 2 mM VPA for 48 h, although a trend to increased expression is suggested in the latter (Fig. 4 A). As regards *DNMT3B*, a significant increase in mRNA expression levels was detected for cells treated with 2 mM VPA for 48 h (Fig. 4 B). When considering WB results, no significant changes at p < 0.05 level were detected for the abundance of DNMT1 and DNMT3B proteins following VPA treatments (Fig. 4 C, D). In the case of DNMT3B, a trend to increased protein abundance values after 2 mM VPA for 48 h is suggested; a statistical significance could not be demonstrated in this case probably because of a large data variability.

**Fig. 4.**
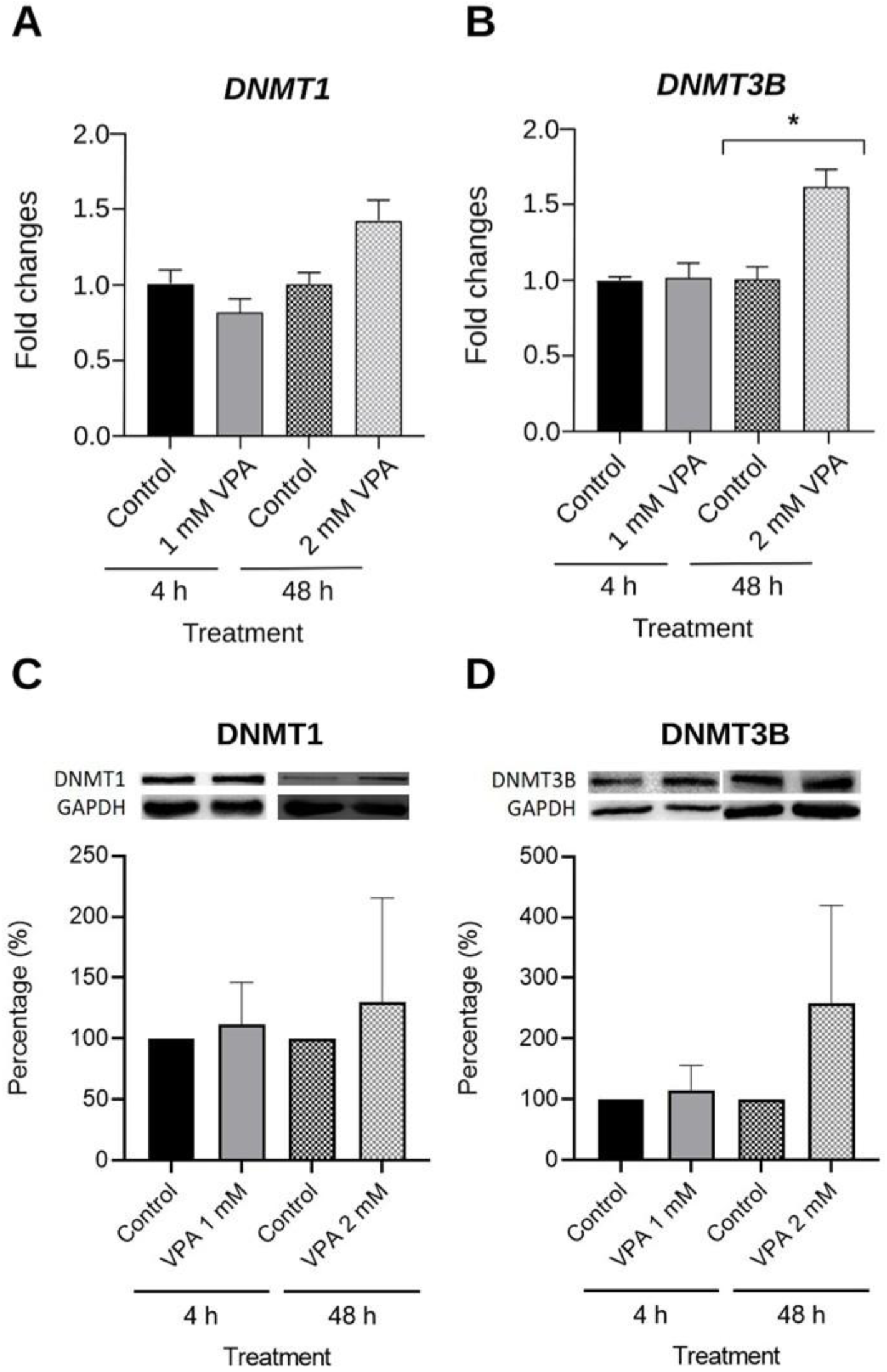
DNMT1 and DNMT3B gene expression levels and protein abundance in VPA-treated HeLa cells as assessed using RT-qPCR (A, B) and WB (C, D). mRNA expression levels of the *DNMT1* and *DNMT3B* genes normalized to endogenous GAPDH control showed a significant increase for *DNMT3B* and a trend toward increase for *DNMT1* after cell treatment with 2 mM VPA for 48 h (A, B). WB and respective densitometry did not indicate statistically significant changes in DNMT1 and DNMT3B protein abundance following VPA treatments, although a trend toward increased values is observed for DNMT3B (C, D). The vertical lines above the graphical bars represent the standard deviation of the mean. Housekeeping GAPDH was used as a loading control.

## 4. Discussion

The present results indicate that the increased frequency of the advanced stage of mitotic catastrophes – those addressed to apoptosis – occurred in HeLa cells treated with 5 mM VPA for 48 h. Specifically, the less packed multinucleated bodies contained within the cellular structures became deeply condensed. Simultaneously, this phenomenon was accompanied by a significant decrease in mitotic chromosome abnormalities. A link between chromosome segregation disorders and mitotic catastrophe has been previously reported (Sazonova et al., 2021).

Although the frequency of images showing typical apoptotic vesicles freely distributed in the middle of non-apoptotic cells, revealing occurrence of apoptosis not properly associated with mitotic catastrophe, was induced by a milder VPA treatment, it was not reflected in the frequency of chromosome abnormalities, and it was shown here only as a complement of the investigation of mitotic catastrophe in VPA-induced cell death.

Cells treated with 1 mM VPA for 4 h did not exhibit changes in the intensity of fluorescent signals indicative of caspase-2, a marker of mitotic catastrophe (Castedo et al., 2004; Vitale et al., 2017; Sazonova et al. 2021). In contrast, a significant enhancement of these signals occurred under 20 mM VPA treatment for 4 or 48 h. This finding is consistent with the morphological images and indicates that mitotic catastrophe can be induced even under a short exposure to VPA at elevated doses. Caspase-2 has been proposed to eliminate non-diploid cells, thereby preserving karyotype stability (Dawar et al., 2016; Vitale et al., 2017). Although the involvement of apoptosis mediated by caspases-3, -8, and -9 in HeLa cells under VPA treatment have been well established (Chen et al., 2006; Hen et al., 2013), the observation of increased caspase-2 levels associated with mitotic catastrophe under the same treatment is reported here for the first time.

Because the p53 effector has been reported to be linked to caspase-2 activation during mitotic catastrophe events (Castedo et al., 2004; Vitale et al., 2017), its incidence was evaluated in HeLa cells following VPA treatments. The results showed a trend toward increased fluorescence intensity of p53 signals after 20 mM VPA treatment for 48 h, but not after 4 h. This was supported by reports indicating that a significant upregulation of p53 expression in HeLa cells, as detected by RT-PCR, occurs only after 24 or 48 h of treatment with elevated VPA concentrations (25 mM and 50 mM) (Hashemi et al., 2022; Kar et al., 2026). Moreover, no effect on p53 expression has been reported in HeLa cells after acute treatment with 1.2, 2.4, and 5.0 mM VPA for 72 h (Sami et al., 2008). However, since caspase-2 fluorescence signals increased significantly after both 20 mM VPA for 48 h and for 4 h, this may suggest that p53-independent effector pathways act during the short time treatment period, in which caspase-2 limits chromosomal instability (Vitale et al., 2017). Degradation of p53 by E6 protein expression - assumed to occur in HeLa cells naturally infected with HPV - is unlikely, since VPA has been reported to induce p53 hyperacetylation, protecting it from E6-mediated degradation (de la Cruz-Hernández et al., 2007).

Although the incidence of mitotic catastrophe increased in HeLa cells under VPA treatment, this effect was not accompanied by decreased DNMT1 levels. This differs from reports in other human tumor models where an increased incidence of mitotic catastrophe was associated with DNMT1 depletion (Chen et al., 2007). The effect of prolonged VPA treatment increasing total and mitochondrial DNMT1 has also been reported in mouse 3T3 cells (Chen et al., 2012). Regarding the expression levels and protein abundance of DNMT3B, assessed by RT-qPCR and Western blotting, they were also not reduced following VPA treatment. DNMT3B expression, in particular, increased significantly after treatment with 2 mM VPA for 48 h.

DNMT1 and DNMT3B have distinct roles and mechanisms of action. Both of them have been identified in several cell types, including HeLa cells (Geiman et al., 2004; Zhang et al., 2011; Rocha et al., 2019). DNMT1 primarily maintains DNA methylation status during DNA replication, whereas DNMT3B establishes new methylation patterns by adding methyl groups to unmethylated cytosines in the genome, thereby creating de novo methylation marks that can influence gene expression (Lyko, 2018). DNMT3B has been found to co-localize with the condensin complex, suggesting a role in mitotic chromosomal events (Geiman et al., 2004). The coexistence of DNMT1 and DNMT3B has been suggested to help maintain genome and epigenome integrity (Hervouet et al., 2018).

Considering that chromatin decondensation occurs in HeLa interphase cells upon histone deacetylase inhibition - accompanied by histone acetylation induced by VPA (Felisbino et al., 2011; Mello, 2021) - the access of transcription factors to promoter regions of the DNMT genes may be facilitated. Additionally, because the length of the spacing between nucleosomes has recently been reported to be crucial for regulating DNMT3B binding to DNA CpG methylation (Xie et al., 2026), it is possible that the increased DNMT3B production while favored by chromatin remodeling enhances specific DNA CpG binding.

### Conclusion

The present study revealed the capability of VPA to induce metabolic pathways addressing to different forms of cell death in HeLa cells. VPA, a drug which is well known to induce apoptosis, was also demonstrated to be capable to induce increased levels of mitotic catastrophe concomitant with reduction of mitotic chromosome abnormalities, thus revealing its oncosuppresive potential. Mitotic catastrophe was identified not only morphologically but also by increased induction of caspase-2, and involvement of p53, at least under more drastic VPA treatment. The incidence of this cell death form was not found to be associated to decreased DNMT1 or DNMT3B gene expressions or protein abundance under the experimental conditions reported here. The significantly increased expression of DNMT3B that was detected following VPA action - if associated with its protein abundance - may be acillitated by the chromatin decondensation consequent from the action of this drug for access of transcription factors to its gene promoter regions and favor specific DNA CpG binding.

## Supporting information

Beatriz Suppl. Western Blots

## Author contributions statement

M.L.S.M. conceived and designed the experiments. B.P.S., C.B.O., M.M.F., M.G.S., and M.A.R. performed the experiments. B.P.S., M.M.F., M.G.S., and M.L.S.M. analyzed the data. M.L.S.M. contributed the reagents/materials/analysis tools and wrote the original draft of the manuscript. All the authors have read and approved the final version of the manuscript.

## Acknowledgments

This research was supported by Fundação de Amparo à Pesquisa do Estado de São Paulo (FAPESP, Brazil) and Conselho Nacional de Desenvolvimento Científico e Tecnológico (CNPq, Brazil). The funders played no role in the study design, data collection and analysis, decision to publish, or preparation of the manuscript. B.P.S. received a fellowship from FAPESP (grant no. 2024/00772-8). M.M.F., M.G.S. and M.L.S.M. received fellowships from CNPq (grants no. 105467/2022-7, 105626/2022-8, and 304589/2024-1, respectively). M.A.R. was recipient of a fellowship from the Coordenação de Aperfeiçoamento de Pessoal de Nível Superior (CAPES, Brazil; Finance code 001). The authors thank Dr. Roger Castilho and Dr. Ana Paula Dalla Costa, from the Department of Clinical Pathology, Faculty of Medicine, UNICAMP, Dr. Sílvio R. Consonni, from the Department of Biochemistry and Tissue Biology, Institute of Biology, UNICAMP, for help with development of caspase and p53 fluorescence protocols, and Mr. Eli H.M. dos Anjos, for his technical assistance. The authors also thank the Obesity and Comorbidities Research Center (Institute of Biology, UNICAMP) and the Central Laboratory of High-Performance Technologies in Life Sciences (LaCTAD, UNICAMP) for providing them with technical facilities.

## Declaration of generative AI and AI-assisted technologies

ChatGPT was used to revise grammar and style language of the manuscript.

## References

Castedo, M., Perfettini, J.L., Roumier, T., Andreau, K., Medema, R., Kroemer, G., 2004. Cell death by mitotic catastrophe: a molecular definition. Oncogene 12, 2825–2837. 10.1038/sj.onc.1207528

Chen, H., Dzitoyeva, S., Manev, H., 2012. Effect of valproic acid on mitochondrial epigenetics. Eur. J. Pharmacol. 5, 51–59. 10.1016/j.ejphar.2012.06.019

Chen, J., Ghazawi, F.M., Bakkar, W., Li, Q., 2006. Valproic acid and butyrate induce apoptosis in human cancer through inhibition of gene expression of Akt/protein kinase K. Mol. Cancer 5, 71. 10.1186/1476-4598-5-71

Chen, T., Hevi, S., Gay, F., Tsujimoto, N., He, T., Zhang, B., Ueda, Y., Li, E., 2007. Complete inactivation of DNMT1 leads to mitotic catastrophe in human cancer cells. Nature Genet. 39, 391–396. 10.1038/ng1982

Dawar, S., Lim, Y., Puccini, J., White, M., Thomas, P., Bouchier-Hayes, L., Green, D.R., Dorstyn, L., Kumar, S., 2017. Caspase-2-mediated cell death is required for deleting aneuploid cells. Oncogene 36, 2704–2714. 10.1038/onc.2016.423

Dejligbjerg, M., Grauslund, M., Litman, T., Collins, L., Qian, X., Jeffers, M., Lichenstein, H., Jenssen, P.B., Sehested, M., 2008. Differential effects of class I isoform histone deacetylase depletion and enzymatic inhibition by belinostat or valproic acid in HeLa cells. Mol. Cancer 7, 70. 10.1186/1476-4598-7-70

de la Cruz-Hernández, E., Pérez-Cárdenas, E., Contreras-Paredes, A., Cantú, D., Mohar, A., Lizano, M., et al., 2007. The effects of DNA methylation and histone deacetylase inhibitors on human papillomavirus early gene expression in cervical cancer, an in vitro and clinical study. Virol. J. 4, 18. 10.1186/1743-422X-4-18

Eyal, S., Yagen, B., Sobol, E., Altschuler, Y., Shmuel, M., Bialer, M., 2004. The activity of antiepileptic drugs as histone deacetylase inhibitors. Epilepsia 45, 737–744. 10.1111/j.0013-9580.2004.00104.x

Felisbino, M.B., Tamashiro, W.M.S.C., Mello, M.L.S., 2011. Chromatin remodeling, cell proliferation and cell death in valproic acid-treated HeLa cells. PLoS ONE 6, e29144. 10.1371/journal.pone.0029144

Galluzzi, L., Vitale, I., Abrams, J.M., Alnemri, E.S., Baehrecke, E.H., Blagosklonny, M.V., Dawson, T.M., Dawson, V.L., El-Deiry, W.S., Fulda, S., Gottlieb, E., Green, D.R., Hegartner, M.O., Kepp, O., Knight, R.A., Kumar, S., Lipton, S.A., Lu, X., Madeo, F., Malorni, W., Mehlen, P., Nuñez, G., Peter, M.E., Piacentini, M., Rubinsztein, D.C., Shi, Y., Simon, H-U., Vandenabeele, P., White, E., Yuan, J., Zhivotovsky, B., Melino, G., Kroemer, G., 2012. Molecular definitions of the Nomenclature Committee on Cell Death 2012. Cell Death Differ. 19, 107–120. 10.1038/cdd.2011.96

Geiman, T.M., Sankpal, U.T., Robertson, A.K., Chen, Y., Mazumdar, M., Heale, J.T., Schmiesing, J.A., Kim, W., Yokomori, K., Zhao, Y., Robertson, K.D., 2004. Isolation and characterization of a novel DNA methyltransferase complex linking DNMT3B with components of the mitotic chromosome condensation machinery. Nucleic Acids Res. 32, 2716–2729. 10.1093/nar/gkh589

Göttlicher, M., Minucci, S., Zhu, P., Kramer, O.H., Schimpf, A., Giavara, S., Sleeman, J.P., LoCoco, F., Nervi, C., Pelicci, P.G., Heinzel, T., 2001. Valproic acid defines a novel class of HDAC inhibitors inducing differentiation of transformed cells. EMBO J. 20, 6969–6978. 10.1093/emboj/20.24.6969

Han, B.R., You, B.R., Park, WHan, B.R., You, B.R., Park, W.H., 2013. Valproic acid inhibits the growth of HeLa cervical cancer cells via caspase-dependent apoptosis. Oncol. Rep. 30, 2999–3005. 10.3892/or.2013.2747

Hashemi, N., Zoshk, M.Y., Rashidian, A., Laripour, R., Fasihi, H., Hami, Z., Chamanara, M., 2022. Anti-proliferative and apoptotic effects of valproic acid on HeLa cells. Int. J. Cancer Manag. 15, e120224. 10.5812/ijcm-120224

Hervouet, E., Peixoto, P., Delage-Mourroux, R., Boyer-Guittaut, M., Cartron, P.F., 2018. Specific or not specific recruitment of DNMTs for DNA methylation, an epigenetic dilema. Clin. Epigenet. 10, 17. 10.1186/s13148-018-0450-y

Kar, R., Almeida, E.A., Basak, D., Roy, A., 2026. Effect of valproic acid on expression of apoptotic markers in HeLa cell line – An *in vitro* study. Int. J. Mol. Immun. Oncol. 11, 48–54. 10.25259/IJMIO_36_2025

Livak, K.J., Schmittgen, T.D., 2001. Analysis of relative gene expression data using real-time quantitative PCR and the 2-(DeltaDeltaC(T)) method. Methods 25, 402–408. 10.1006/meth.2001.1262

Lyko, F., 2018. The DNA methyltransferase family: a versatile toolkit for epigenetic regulation. Nature Rev. Genet. 19, 81–92. 10.1038/nrg.2017.80

Mello, M.L.S., 2021. Sodium valproate-induced chromatin remodeling. Front. Cell Dev. Biol. 9, 645518. 10.3389/fcell.2021.645518

Mello, M.L.S., Vidal, B.C., 2017. The Feulgen reaction: a brief review and new perspectives. Acta Histochem. 119, 603–609. 10.1016/j.acthis.2017.07.002

Miranda, E.I., Santana, C., Rojas, E., Hernández, S., Ostrosky-Wegman, P., García-Carrancá, A., 1996. Induced mitotic death of HeLa cells by abnormal expression of c-H-*ras*. Mut. Res. 349, 173–182.

Peng, D.F., Kanai, Y., Sawada, M., Ushijima, S., Hiraoka, N. Ktazawa, S., Hirohashi, S., 2006. DNA methylation of multiple tumor-related genes in association with overexpression of DNA methyltransferase 1 (DNMT1) during multistage carcinogenesis of the pancreas. Carcinogenesis 27, 1160–1168. 10.1093/carcin/bgi361

Phiel, C.J., Zhang, F., Huang, E.Y., Guenther, M.G., Lazar, M.A., Klein, P.S., 2001. Histone deacetylase is a direct target of valproic acid, a potent anticonvulsant, mood stabilizer, and teratogene. J. Biol. Chem 276, 36734–36741. 10.1074/jbc.M101287200

Rocha, M.A., Oliveira, C.B., Mello, M.L.S., 2021. Sodium valproate cytotoxicity effects as assessed by the MTT assay. Research Data Repository, UNICAMP– versions 1–3. 10.25824/redu/XPTX4F

Rocha, M.A., Veronezi, G.M.B., Felisbino, M.B., Gatti, M.S.V., Tamashiro, W.M.S.C., Mello, M.L.S., 2019. Sodium valproate and 5-aza-2’-deoxycytidine differentially modulate DNA demethylation in G1 phase-arrested and proliferative HeLa cells. Sci. Rep. 9, 18236. 10.1038/s41598-019-54848-x

Rocha, M.A., Vidal, B.C., Mello, M.L.S., 2023. Sodium valproate modulates the methylation status of lysine residues 4, 9 and 27 in histone H3 of HeLa cells. Curr. Mol. Pharmacol. 16, 197–210. 10.2174/1874467215666220316110405

Romoli, M., Mazzocchetti, P., D’Alonzo, R., Siliquini, S., Rinddi, V.E., Verrotti, A., Calabresi, P., Costa, C., 2019. Valproic acid and epilepsy: from molecular mechanisms to clinical evidences. Curr. Neuropharmacol. 17, 926–946. 10.2174/1570159X17666181227165722

Saito, Y., Kanai, Y., Nakagawa, T., Sakamoto, M., Saito, H., Ishii, H., Hirohashi, S., 2003. Increased protein expression of DNA methyltransferase (DNMT) 1 is significantly correlated with the malignant potential and poor prognosis of human hepatocellular carcinomas. Int. J. Cancer 105, 527–532. 10.1002/ijc.11127

Sami, S., Höti, N., Xu, H.M., Shen, Z., Huang, X., 2008. Valproic acid inhibits the growth of cervical cancer both *in vitro* and *in vivo*. J. Biochem. 144, 357–362. 10.1093/jb/mvn074

Sargolzaei, J., Rabbani-Chadegani, A., Mollaei, H., and Deezagi, A., 2017. Spectroscopic analysis of the interaction of valproic acid with histone H1 in solution and in chromatin structure. Int. J. Biol. Macromol. 99, 427–432. 10.1016/j.ijbiomac.2017.02.098

Sazonova, E., Petrichuk, S., Kopeina, G.S., Zhivotovsky, B., 2021. A link between mitotic defects and mitotic catastrophe: detection and cell fate. Biol. Direct 16, 25. 10.1186/s13062-021-00313-7

Sforça, B.P., Mello, M.L.S., 2026. Markers for mitotic catastrophe in sodium valproate-treated HeLa cells. 10.25824/redu/8OEBZ7, Repository of Research Data, UNICAMP, V1, UNF:6:m8UAxOJ1Ci8GhliXj0R1A==[fileUNF]

Sforça, B.P., Furtado, M.M., Santos, M.G., Mello, M.L.S., 2026. Mitotic defects and cell death in sodium valproate-treated HeLa cells. 10.25824/redu/55WSVA, Repository of Research Data, UNICAMP, V1, UNF:6:8ZORtVBnqFyoBuHzqGK9mA==[fileUNF]

Swanson, P.E., Carroll, S.B., Zhang, X.F., Mackey, M.A., 1995. Spontaneous premature chromosome condensation, micronucleus formation, and non-apoptotic cell death in heated HeLa S3 cells. Ultrastructural observations. Am. J. Pathol. 146, 963–971. https://www.ncbi.nlm.nih.gov/pubmed/7717463

Veronezi, G.M.B., Felisbino, M.B., Gatti, M.S.V., Mello, M.L.S., Vidal, B.C., 2017. DNA methylation changes in valproic acid-treated HeLa cells as assessed by image analysis, immunofluorescence and vibrational microspectroscopy. PLoS ONE 12, e0170740. 10.1371/journal.pone.0170740

Vidal, B.C., Mello, M.L.S., 2020. Sodium valproate (VPA) interactions with DNA and histones. Int. J. Biol. Macromol. 163, 219–231. 10.1016/j.ijbiomac.2020.06.265

Vitale, I., Galluzzi, L., Castedo, M., Kroemer, G., 2011. Mitotic catastrophe: a mechanism for avoiding genomic instability. Nat. Rev. Mol. Cell Biol. 12, 385–392. 10.1038/nrm3115

Vitale, I., Manic, G., Castedo, M, Kroemer, G., 2017. Caspase 2 in mitotic catastrophe: The terminator of aneuploid and tetraploid cells. Mol. Cell. Oncol. 4, e1299274. 10.1080/23r723556.2017.1299274

Xie, X., Liu, M., Chua, G.N.L., Zhou, E., Dykstra, M., Liu, S., Jones, P.A., Worden, E.J., 2026. Nucleosome spacing regulates linker methylation by DNMT3A2/3B3. Mol. Cell 86, 834–850. 10.1016/j.molcel.2026.01.030

Zhang, W., Xu, J., 2017. DNA methyltransferases and their roles in tumorigenesis. Biomarker Res. 5, 1. 10.1186/s40364-017-0081-z

Zhang, Y., Chen, F.Q., Sun, Y.H., Zhou, S.Y., Li, T.Y., Chen, R., 2011. Effects of DNMT1 silencing on malignant phenotype and methylated gene expression in cervical cancer cells. J. Expt. Clin. Cancer Res. 30, 98. http://www.jeccr.com/content/30/1/98

