## Supplementary material for "Mitotic catastrophe and other cellular instability events in sodium valproate-treated HeLa cells": Beatriz Suppl. Western Blots

**Supplementary Figures**

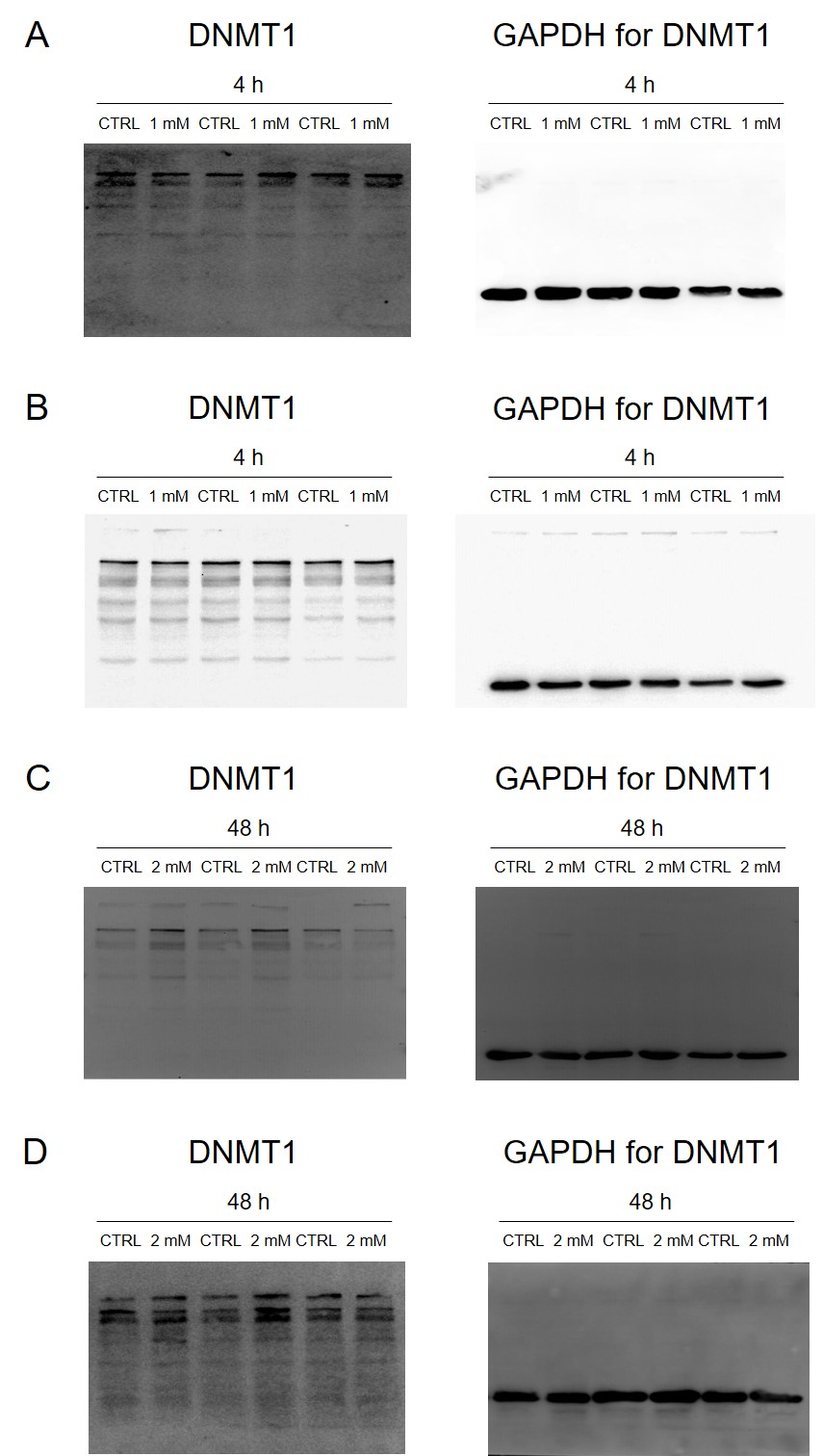

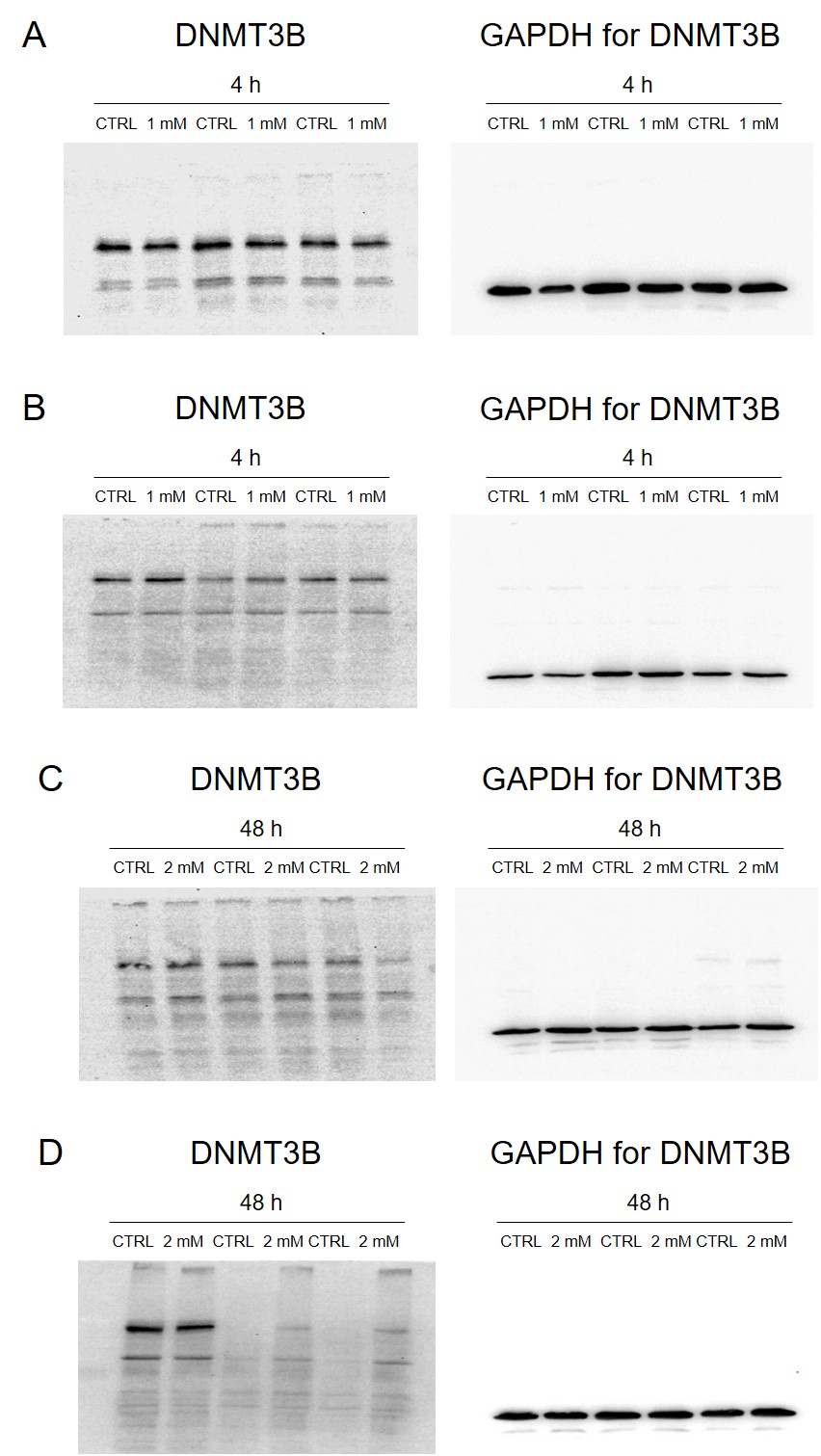
